## Supplementary Information for "SPACA6 structure reveals a conserved superfamily of gamete fusion-associated proteins"

**This PDF file includes:**

Figures S1 to S20
Tables S1 to S2

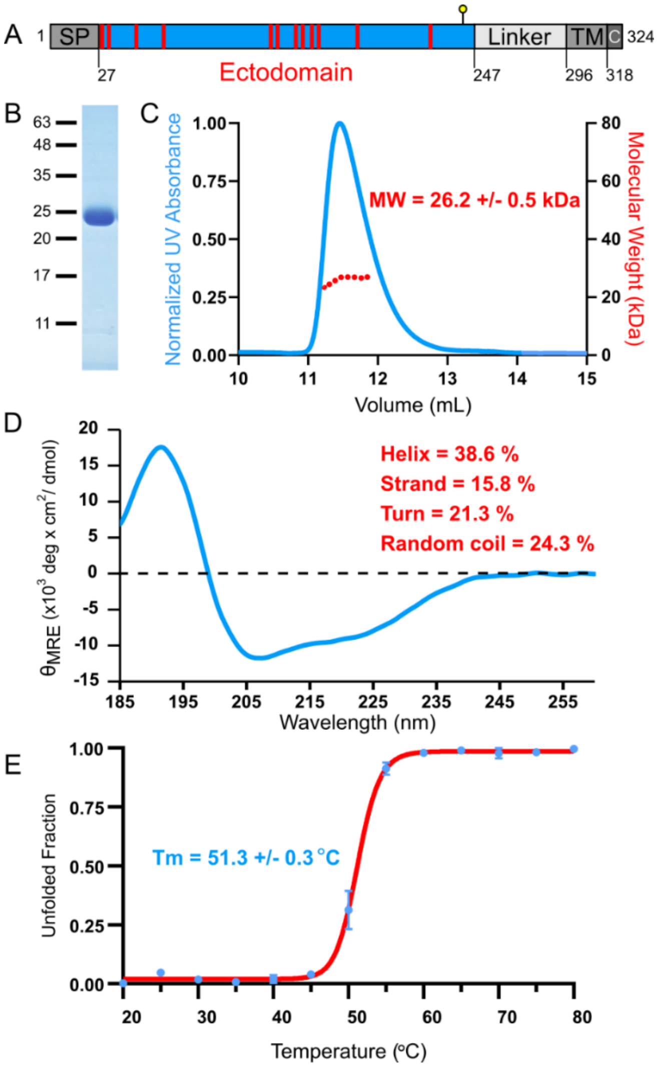

**Fig. S1. SPACA6 ectodomain expresses as a folded, monomeric protein. A)** Domain architecture of full-length SPACA6. The signal peptide (SP), transmembrane helix (TM), and cytoplasmic tail (C) are indicated in the schematic. Red lines indicate cysteine residues, and yellow lollipop indicates predicted N-linked glycosylation site. **B)** Coomassie-stained SDS-PAGE analysis of purified recombinant SPACA6 ectodomain. Molecular weights based on protein ladder bands are shown on the left. **C)** SEC-MALS analysis of recombinant SPACA6 ectodomain. Size-exclusion chromatogram with normalized UV absorbance (280 nm) is shown in the blue curve with calculated molecular weight values derived from light scattering shown in red dots. The presented molecular weight is the average of calculated molecular weights from each individual light-scattering measurements ($n$=108) taken within the peak, ± standard deviation. **D)** Far-UV CD spectra of recombinant SPACA6 ectodomain at 0.16 mg $mL^{-1}$. Spectra is an average of ten accumulations that are buffer corrected and smoothened. **E)** Thermal melt of recombinant SPACA6 ectodomain. Mean residue ellipticity was recorded at 207 nm. Values are averages of independent triplicates ($n$=3) with error bars for standard deviation shown in blue.

>SPACA6_Native
MALLALASAVPSALLALAVFRVPAWACLLCFTTYSERLRICQMFVGMRSPKLEECEEAFTAAFQG
LSDTEINYDERSHLHDTFTQMTHALQELAAAQGSFEVAFPDAAEKMKKVITQLKEAQACIPPCGL
QEFARRFLCSGCYSRVCDLPLDCPVQDVTVTRGDQAMFSCIVNFQLPKEEITYSWKFAGGGLRTQ
DLSYFRDMPRAEGYLARIRPAQLTHRGTFSCVIKQDQRPLARLYFFLNVTGPPPRAETELQASFR
EVLRWAPRDAELIEPWRPSLGELLARPEALTPSNLFLLAVLGALASASATVLAWMFFRWYCSGN

>SPACA6_Recombinant
MKLCILLAVVAFVGLSLGCLLCFTTYSERLRICQMFVGMRSPKLEECEEAFTAAFQGLSDTEINY
DERSHLHDTFTQMTHALQELAAAQGSFEVAFPDAAEKMKKVITQLKEAQACIPPCGLQEFARRFL
CSGCYSRVCDLPLDCPVQDVTVTRGDQAMFSCIVNFQLPKEEITYSWKFAGGGLRTQDLSYFRDM
PRAEGYLARIRPAQLTHRGTFSCVIKQDQRPLARLYFFLNVTGGRLVPRGSHHHHHHHHHH

>SPACA6_Recombinant_Processed
CLLCFTTYSERLRICQMFVGMRSPKLEECEEAFTAAFQGLSDTEINYDERSHLHDTFTQMTHALQ
ELAAAQGSFEVAFPDAAEKMKKVITQLKEAQACIPPCGLQEFARRFLCSGCYSRVCDLPLDCPVQ
DVTVTRGDQAMFSCIVNFQLPKEEITYSWKFAGGGLRTQDLSYFRDMPRAEGYLARIRPAQLTHR
GTFSCVIKQDQRPLARLYFFLNVTGGRLVPR

**Fig. S2.** Sequences of native SPACA6 and recombinant SPACA6 ectodomain. One-letter amino acid sequences for SPACA6 Native, Recombinant, and the Recombinant construct after secretion and tag-removal with thrombin. Orange indicates amino acids of the four-helix bundle; green indicates hinge region amino acids; blue indicates Ig-like domain amino acids; grey indicates native signal peptide; red indicates native transmembrane helix; black indicates non-ectodomain native sequences; grey highlight indicates BiP signal peptide for *Drosophila* S2 cell secretion; green highlight indicates thrombin cleavage site; black highlight indicates 10x His tag; and purple indicates putative N-linked glycosylation site.

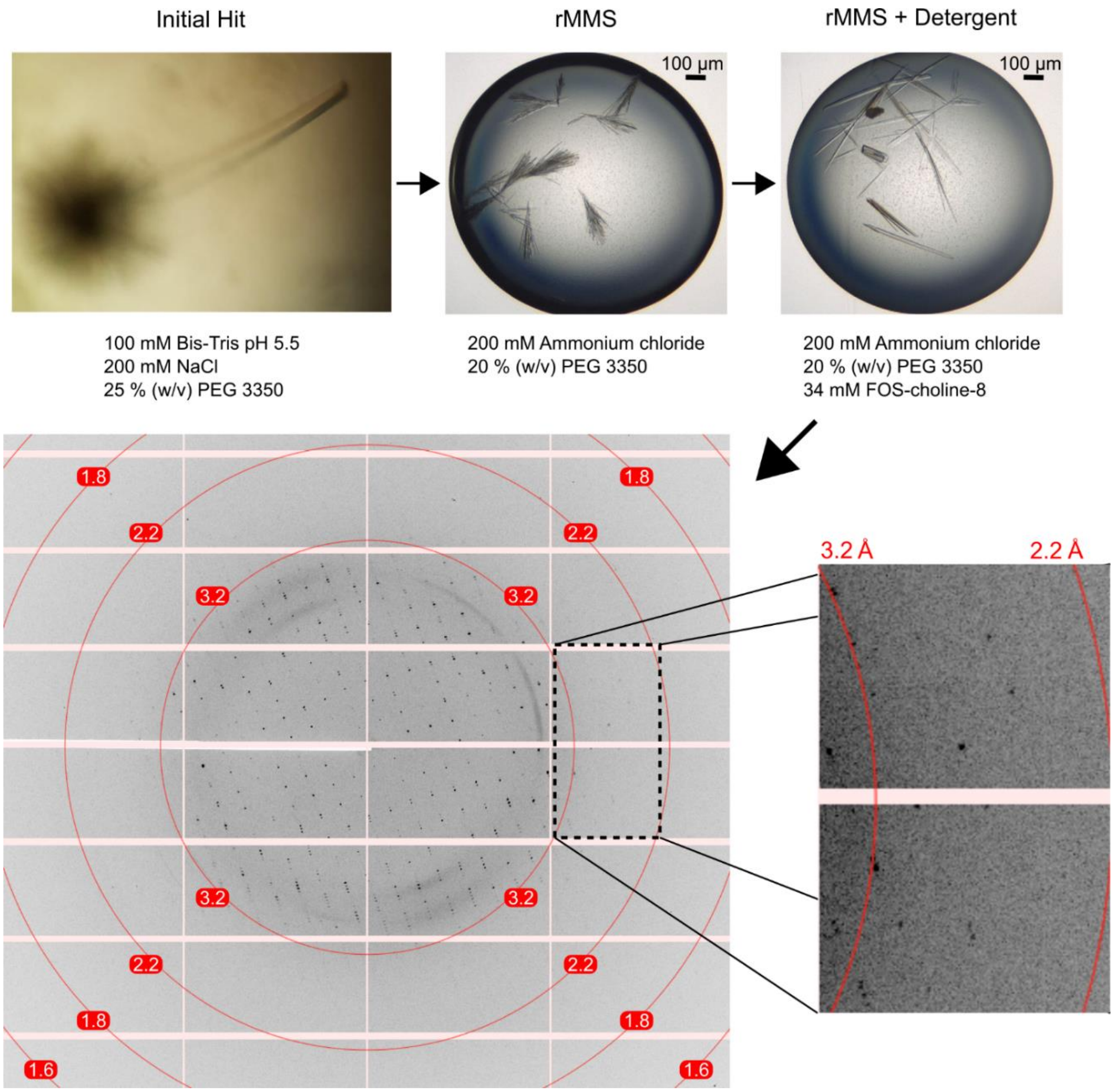

**Fig. S3. Crystal growth and diffraction of SPACA6.** Upper images were taken during optimization; Optimization of SPACA6 crystals (upper images) with the compositions of the reservoir solutions stated the images. Upper left: Initial crystals obtained by vapor-diffusion; these crystals diffracted poorly. Upper middle: Crystals obtained when initial crystals were used as seeds for random microseed matrix screening (rMMS). Upper right: Larger crystals obtained through screening detergents as additives. Lower left: Diffraction image produced from the detergent-stabilized crystals, collected at the NE-CAT (24-ID-C) beamline at the Advanced Photon Source (Argonne National Laboratory, Lemont, IL). Red circles indicate (from innermost to outermost) 3.2 Å, 2.2 Å, 1.8 Å, and 1.6 Å resolution rings. Lower right insets: Higher magnification of boxed region showing high-resolution reflections to 2.2 Å resolution.

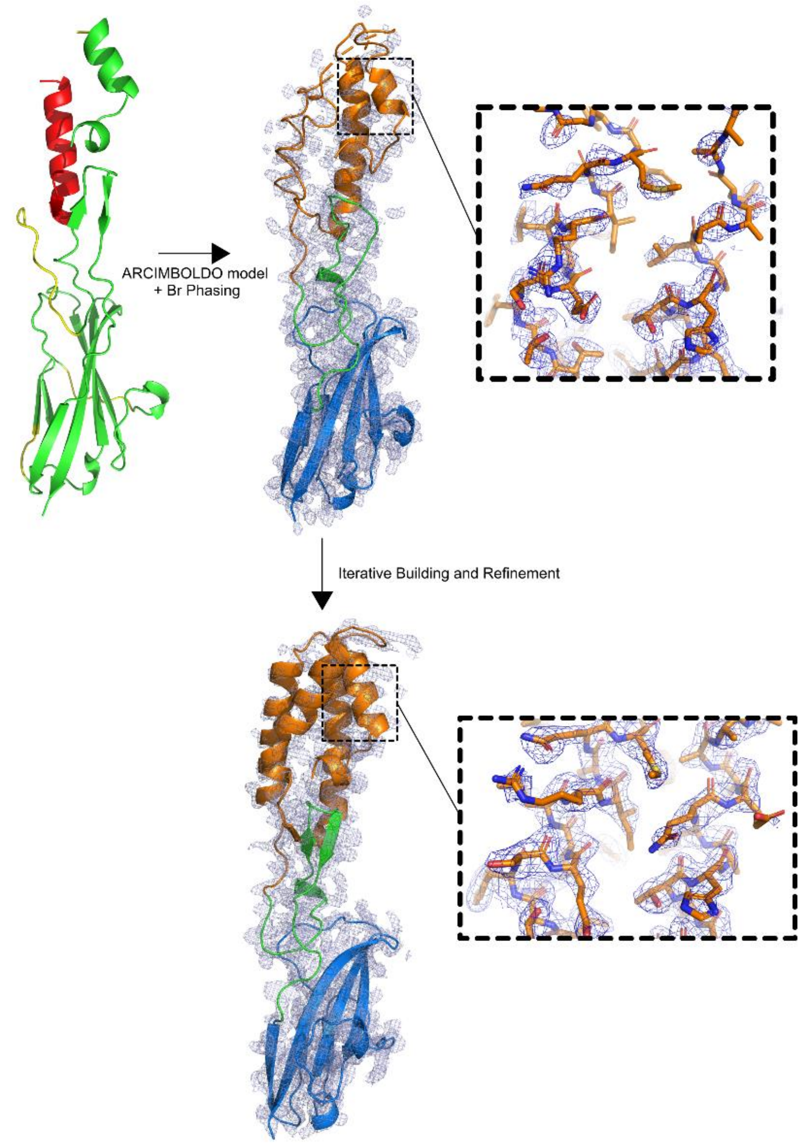

**Fig. S4. Initial ARCIMBOLDO_SHREDDER model and electron density map.** Upper left: Initial SPACA6 model produced via ARCIMBOLDO_SHREDDER (65% complete). Portions of the model built into strong electron density are colored green; portions of the model built into strong electron density but modeled as poly-alanine are colored yellow; and portions of the model unconnected to other parts and made up of poly-alanine are colored red. Upper right: Following Br-SAD phasing and ARCIMBOLDO_SHREDDER molecular replacement, an initial phased combined experimental electron density map ($|F_o|$) map shown at 1$\sigma$ was obtained. The inset shows the poor electron density of the 4HB. Lower: Following iterative building/refinement, the final electron density map ($2|F_o|–|F_c|$) at 1$\sigma$ superimposed with the refined SPACA6 structure is shown. The inset shows the same region of the 4HB as the upper inset. Structures are colored orange for 4HB, green for hinge, and blue for Ig-like domain.

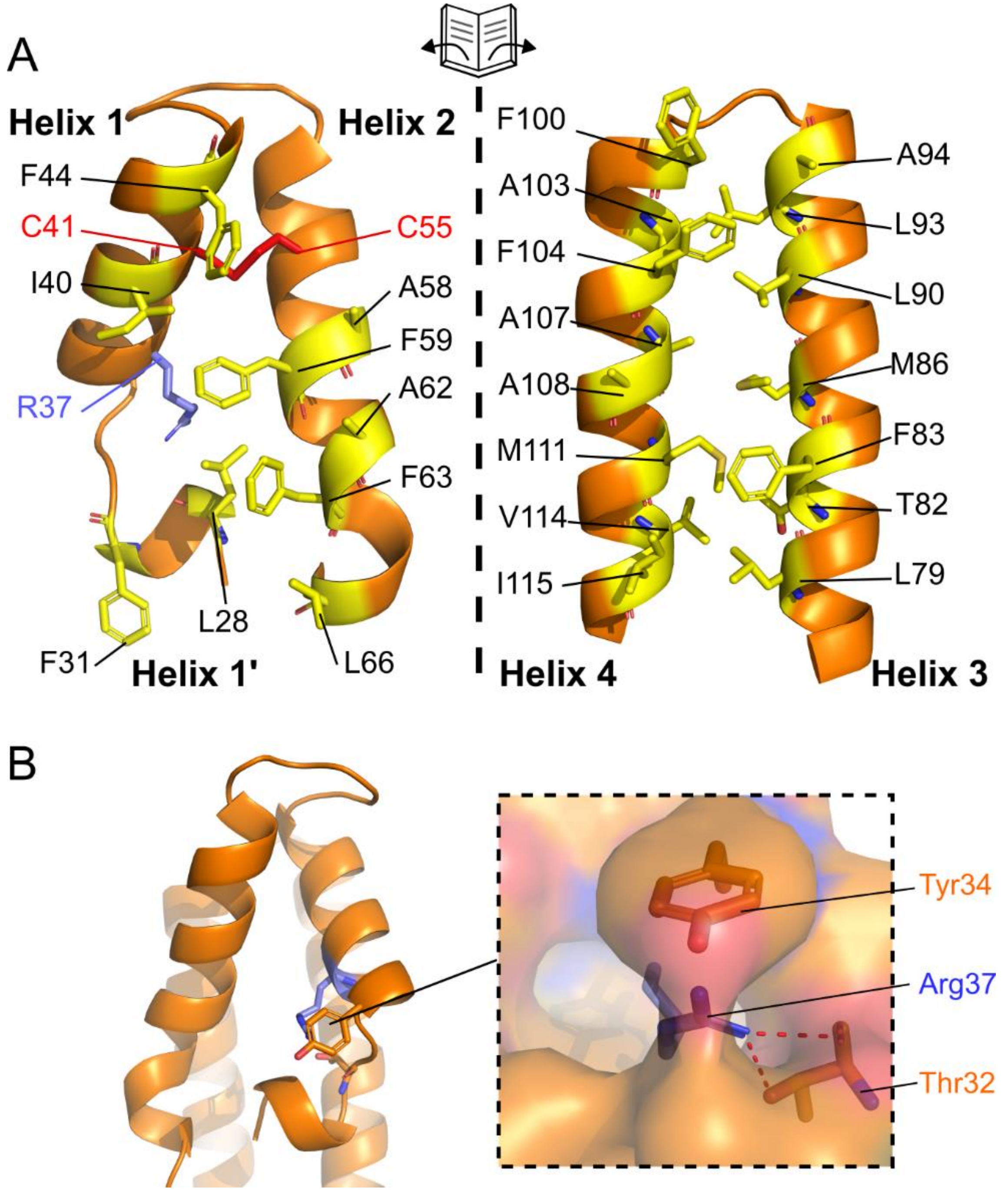

**Fig. S5. Conserved arginine within 4HB hydrophobic core. A)** Cross-section of SPACA6 four-helix bundle. The bundle was split in the middle and opened to reveal the hydrophobic core. Hydrophobic components of the 4HB core are colored yellow and displayed as sticks. Components of the 4HB core include Arg37 (purple) and a Cys41-Cys55 disulfide bond (red). **B)** Triangular face of the 4HB, with Tyr34, Arg37 (purple), and Thr32 shown as sticks. Inset shows the Arg37 interaction with Thr32 and its occlusion by Tyr34.

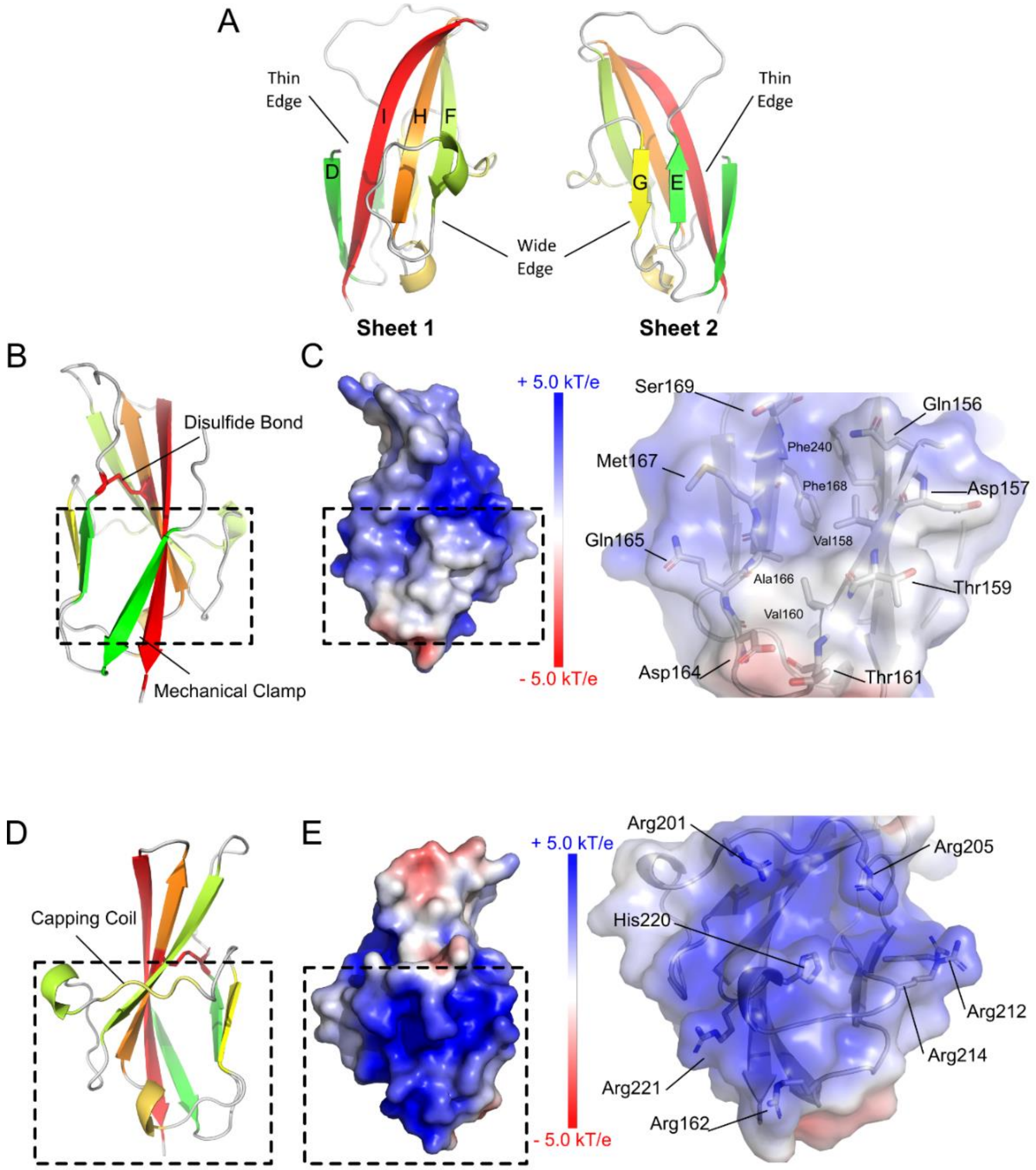

**Fig. S6. C-terminal Ig-like domain of SPACA6. A)** Ribbon diagram of the SPACA6 Ig-like domain with its six strands labelled. **B)** Ribbon diagram of the thinner edge of the Ig-like domain. The disulfide bond is shown as sticks. **C)** Surface electrostatic potential of the thinner edge of the Ig-like domain. Hydrophobic patch outlined by dashed box is shown as a zoomed image (right) with the residues shown as sticks. **D)** Ribbon diagram of the wider edge of the Ig-like domain. Capping coil that covers the hydrogen bonds in Strand F is shown. **E)** Surface electrostatic potential of the wider edge of the Ig-like domain. Positively charged cavity outlined by dashed box is shown as a zoomed image (right) with labeled residues shown as sticks.

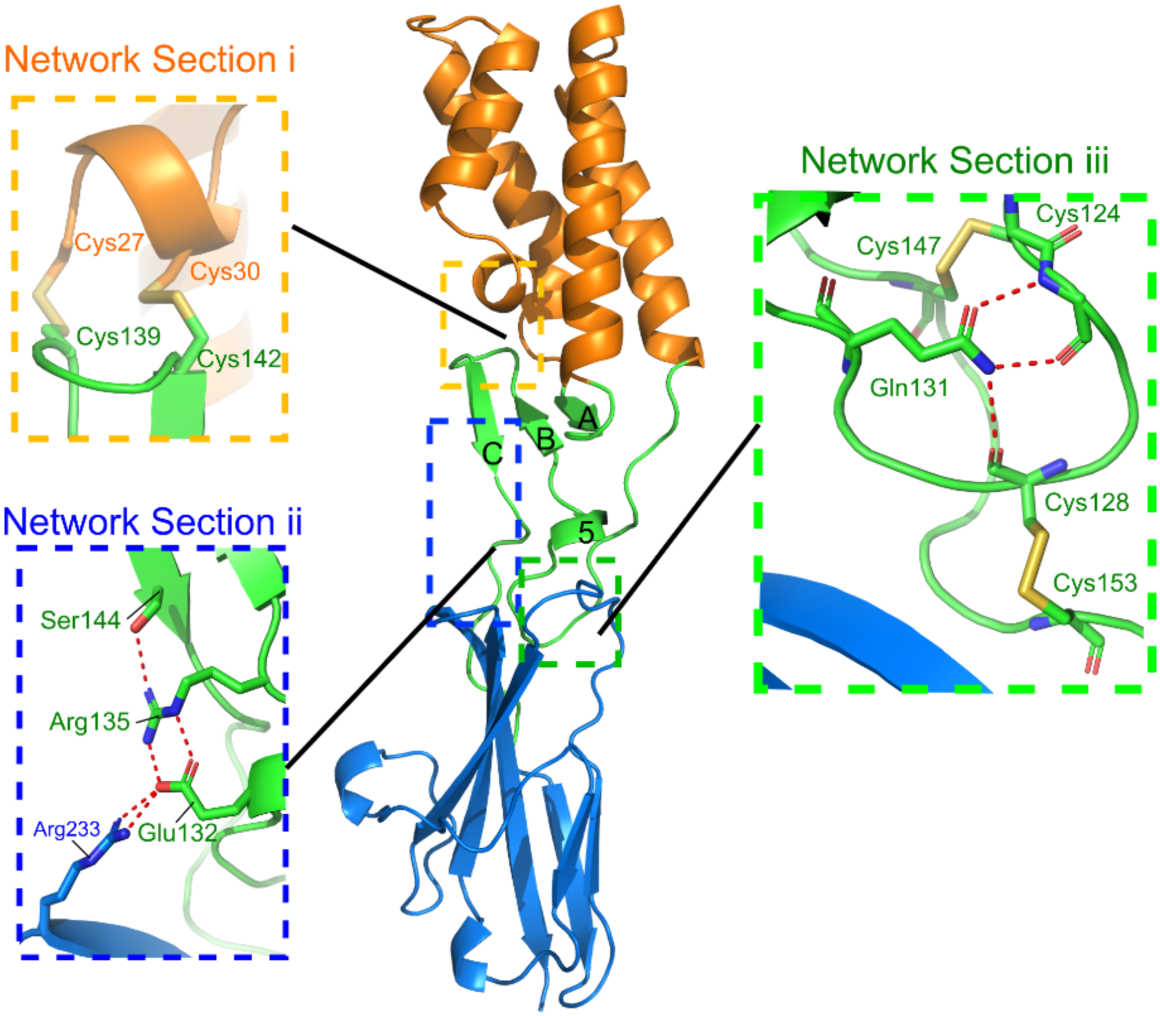

**Fig. S7. SPACA6 hinge region connects the 4HB and Ig-like domains.** Ribbon diagram of the hinge region (green), which connects the 4HB (orange) and the Ig-like domain (blue). The three sections of the covalent/electrostatic network that rigidifies the region are shown in detail.

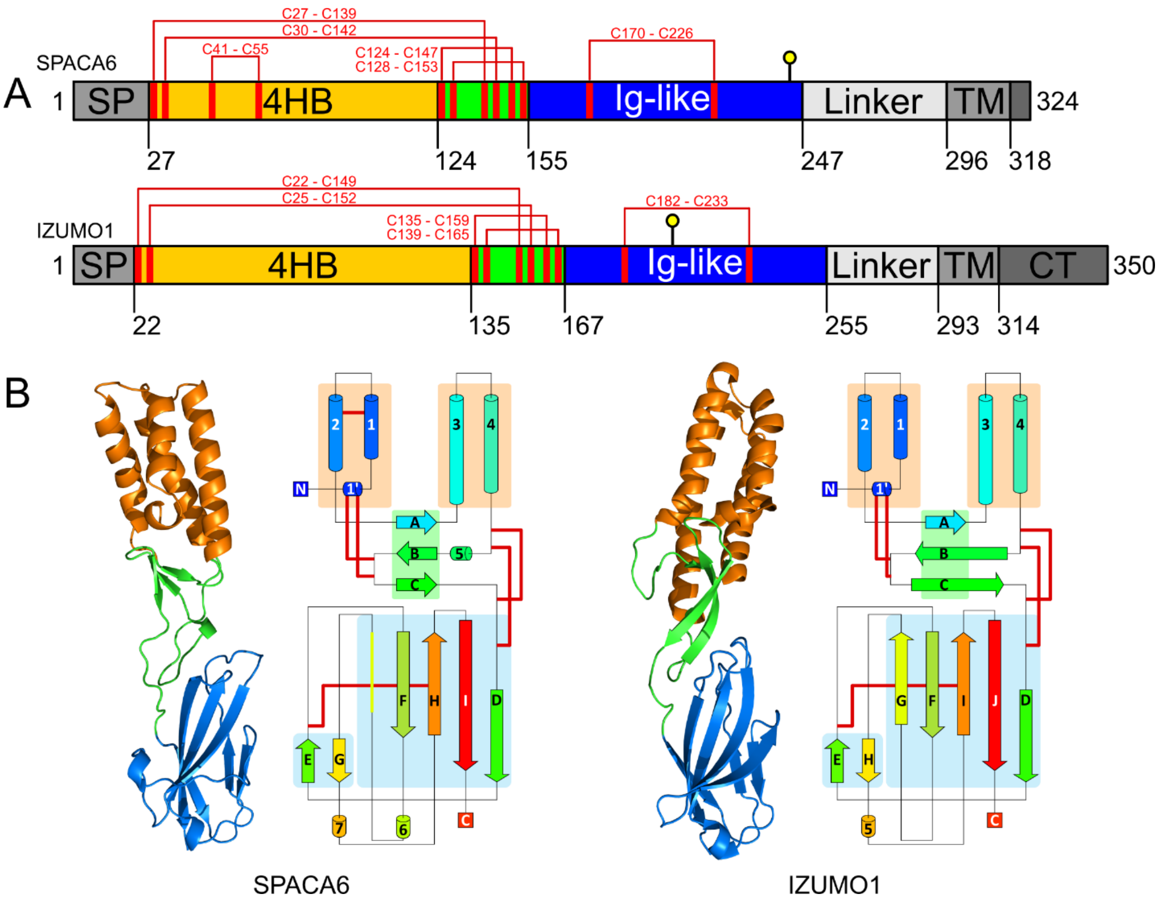

**Fig. S8. SPACA6 and IZUMO1 share a similar structure. A)** Domain architecture schematic of SPACA6 and IZUMO1. Red lines indicate disulfide bonds and yellow lollipop indicates a predicted N-linked glycan site. Abbreviations: SP, signal peptide; TM, transmembrane helix; CT, cytoplasmic tail. **B)** Ribbon and topology diagrams comparing ectodomain structures for SPACA6 and IZUMO1 (PDB: 5F4E). Domains are colored according to panel A.

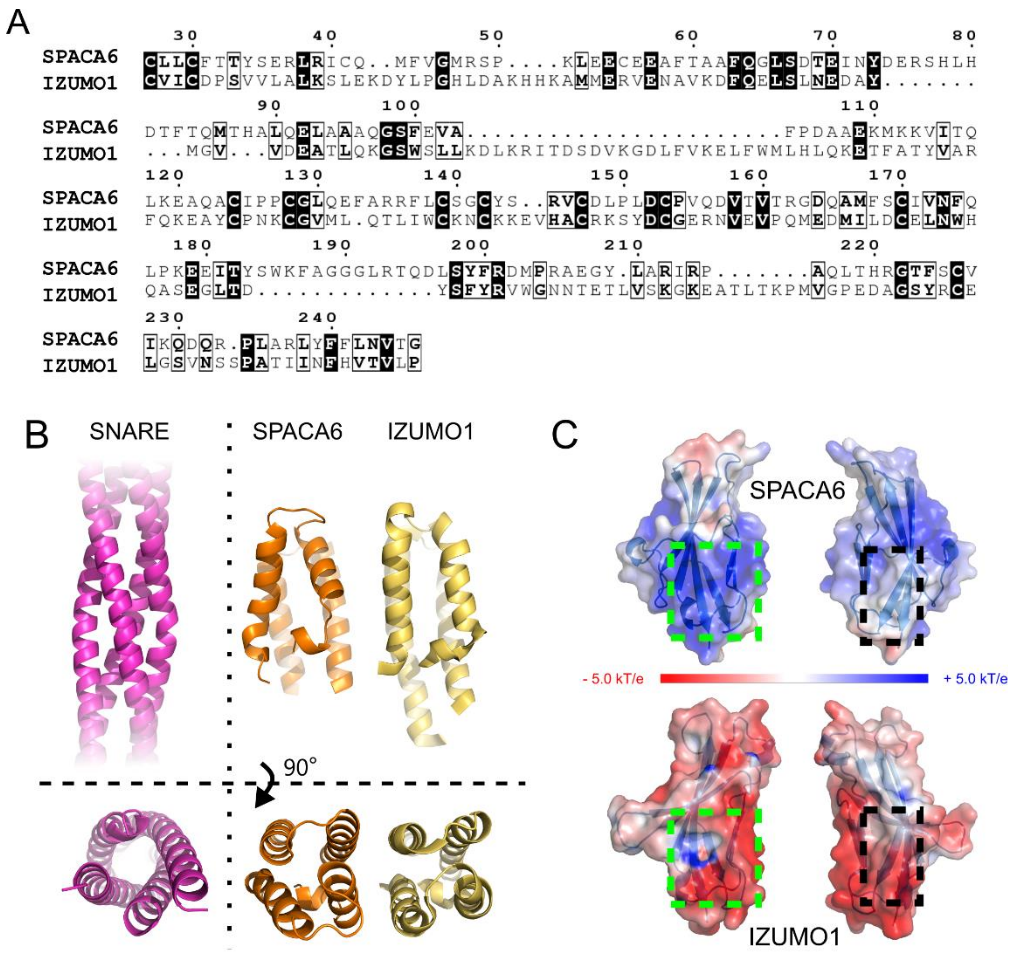

**Fig. S9. Structural comparison of IZUMO1 and SPACA6. A)** Human SPACA6 (RefSeq: NP_001303901) and IZUMO1 (RefSeq: NP_872381) ectodomain sequences aligned using CLUSTAL OMEGA. Strictly conserved residues are displayed as white text on a black background. Residues with conservative substitutions are in bold. **B)** Four-helix bundle comparison in two orientations. Ribbon diagrams for autophagic SNARE complex (PDB: 4WY4, magenta), SPACA6 (orange), and IZUMO1 (PDB: 5F4E, yellow). **C)** Surface electrostatic potential of SPACA6 and IZUMO1 Ig-like domains. The positively charged pocket from SPACA6 is outlined with a dashed green line, and the hydrophobic surface from SPACA6 is outlined with a dashed black line.

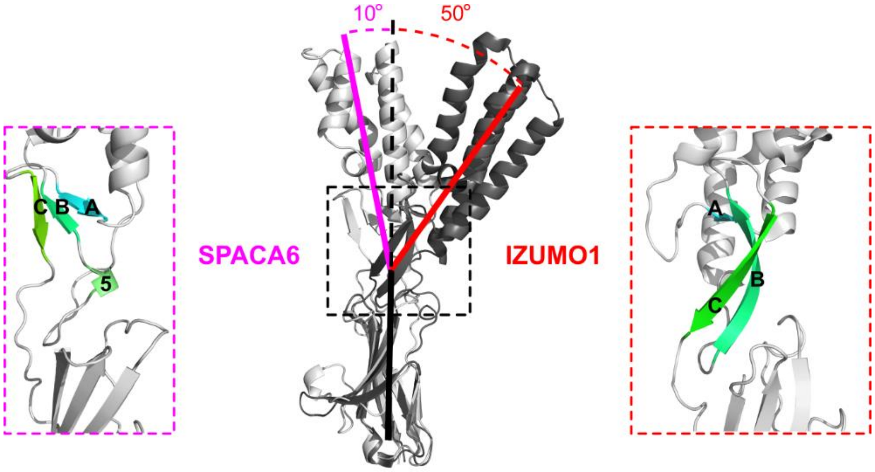

**Fig. S10. Structural evidence of flexibility in IZUMO1 and SPACA6.** SPACA6 (white) and IZUMO1 (PDB: 5F4E, black) structures were aligned by the Ig-like domain to compare the orientation of the 4HB. Insets show the hinge region for both SPACA6 (purple box) and IZUMO1 (red box).

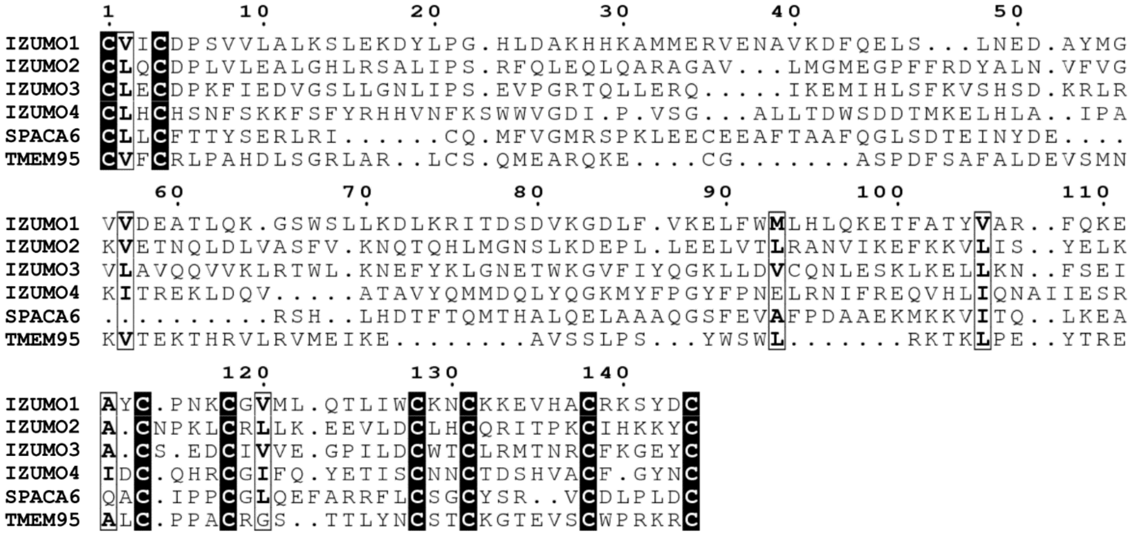

**Fig. S11. Sequence alignment of the 4HB domains from the proposed IST-superfamily.** The 4HB domains from human IZUMO1 (NCBI: NP_872381), IZUMO2 (NCBI: NP_689571), IZUMO3 (NCBI: NP_001351937), IZUMO4 (NCBI: NP_001034935), SPACA6 (NCBI: NP_001303901), and TMEM95 (NCBI: XP_016880054) were aligned using CLUSTAL OMEGA. Strictly conserved residues are displayed as white text on a black background. Residues with conservative substitutions are in bold. Alignment numbering corresponds to the human IZUMO1 sequence.

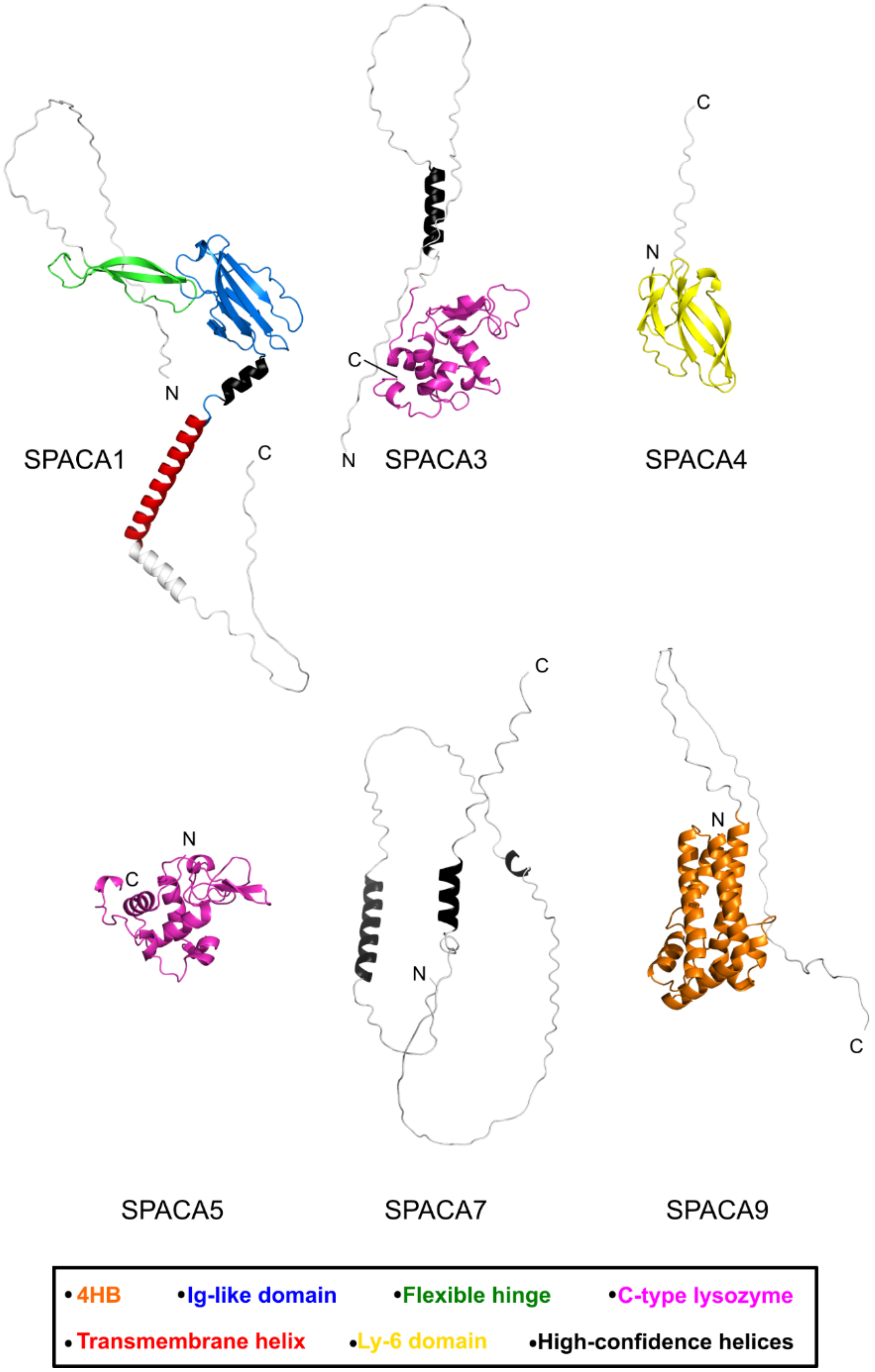

**Fig. S12. AlphaFold predictions of structures of SPACA proteins.** Cartoon representations of AlphaFold structure predictions of human SPACA1 (AF-Q9HBV2-F1), SPACA3 (AF-Q8IXA5-F1), SPACA4 (AF-Q8TDM5-F1), SPACA5 (AF-Q96QH8-F1), SPACA7 (AF-Q96KW9-F1), and SPACA9 (AF-Q96E40-F1). Black helices are non-interacting with high predictive confidence. Areas of low predictive confidence are colored white. Signal peptides predicted for SPACA1, SPACA4, SPACA5, and SPACA7 were removed.

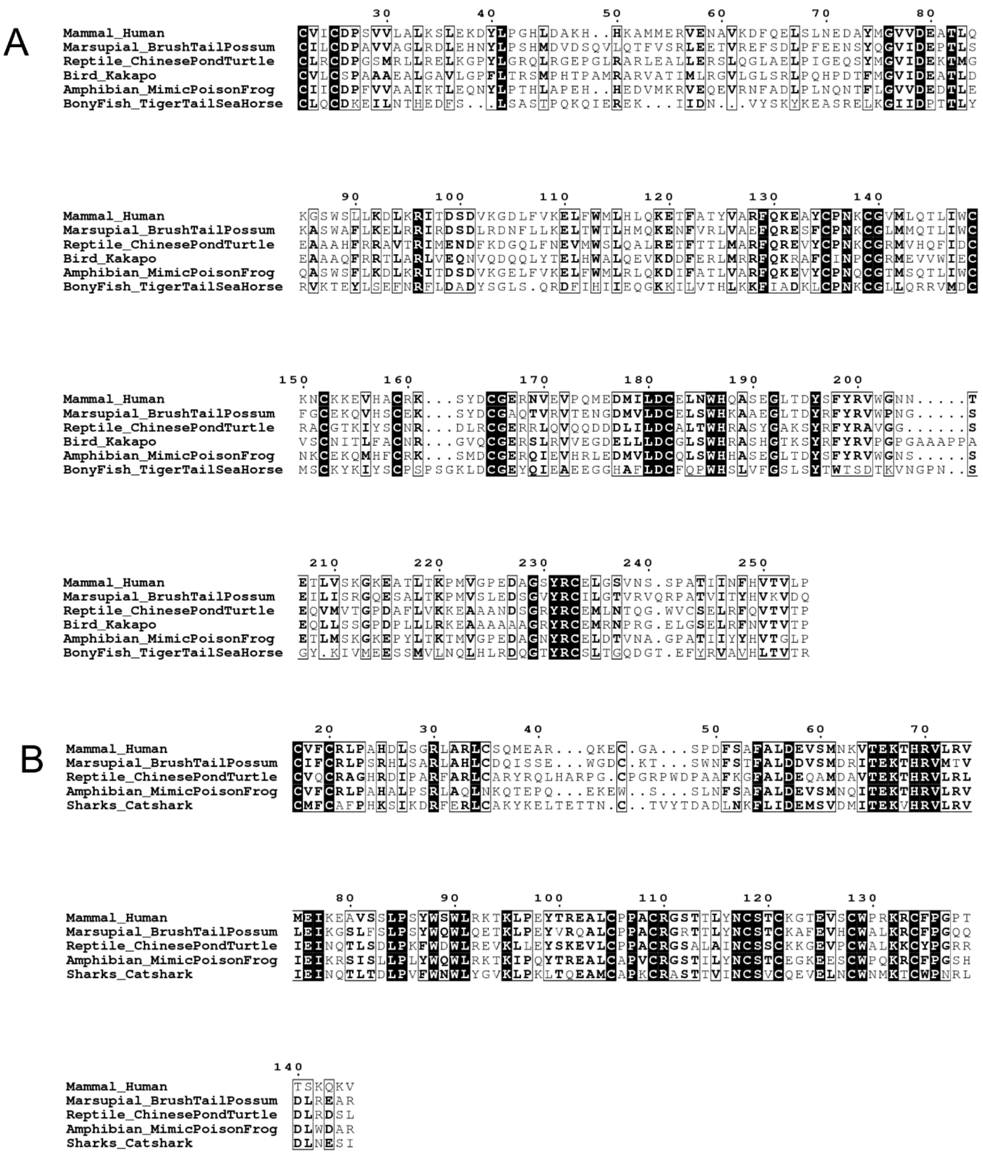

**Fig. S13. Sequence alignment of IZUMO1 and TMEM95 from multiple species. A)** Ectodomain sequences for IZUMO1 from human (*Homo sapiens*, NCBI: NP_872381), the marsupial brush tail possum (*Trichosurus vulpecula,* NCBI: XP_036602972), the Chinese pond turtle (*Mauremys reevesii,* NCBI: XP_039366991), the flightless parrot Kākāpō (*Strigops habroptila*, NCBI: XP_030366609), the mimic poison frog (*Ranitomeya imitator*, NCBI: CAF4957531), and the tiger tail seahorse (*Hippocampus comes*, NCBI XP_019744032) were aligned using CLUSTAL OMEGA. **B)** Ectodomain sequences for TMEM95 from human (*Homo sapiens*, NCBI: XP_016880054), the marsupial brush tail possum (*Trichosurus vulpecula,* NCBI: XP_036622079), the Chinese Pond Turtle (*Mauremys reevesii,* NCBI: XP_039355851), the mimic poison frog (*Ranitomeya imitator*, NCBI: CAF5092145), and the catshark (*Scyliorhinus canicular*, NCBI: XP_038642792) were aligned using CLUSTAL OMEGA. Strictly conserved residues are displayed as white text on a black background. Residues with conservative substitutions are in bold. Both alignments are numbered according to the human sequence.

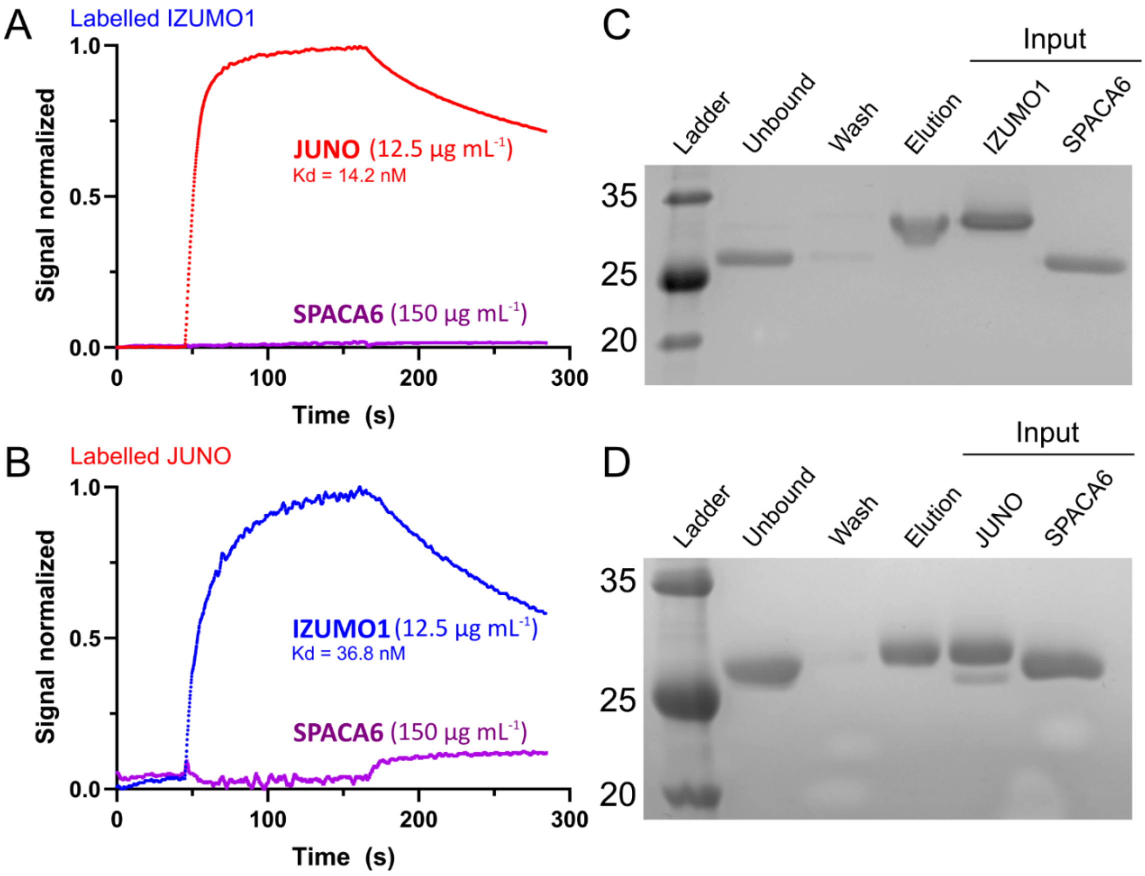

**Fig. S14. SPACA6 does not bind to IZUMO1 and JUNO. A and B)** BLI sensorgrams of biotin-labelled **A)** IZUMO1 or **B)** JUNO as bait with high concentrations of JUNO, IZUMO1 or SPACA6 proteins as the analyte. **C and D)** Pull-down analyses of His-tagged **C)** IZUMO1 or **D)** JUNO as bait with equal concentrations of untagged SPACA6 as analyte protein. SDS-PAGE analyses stained with Coomassie brilliant blue are used to detect the protein. Both BLI and pull-downs results are representative of independent duplicate (*n*=2) studies.

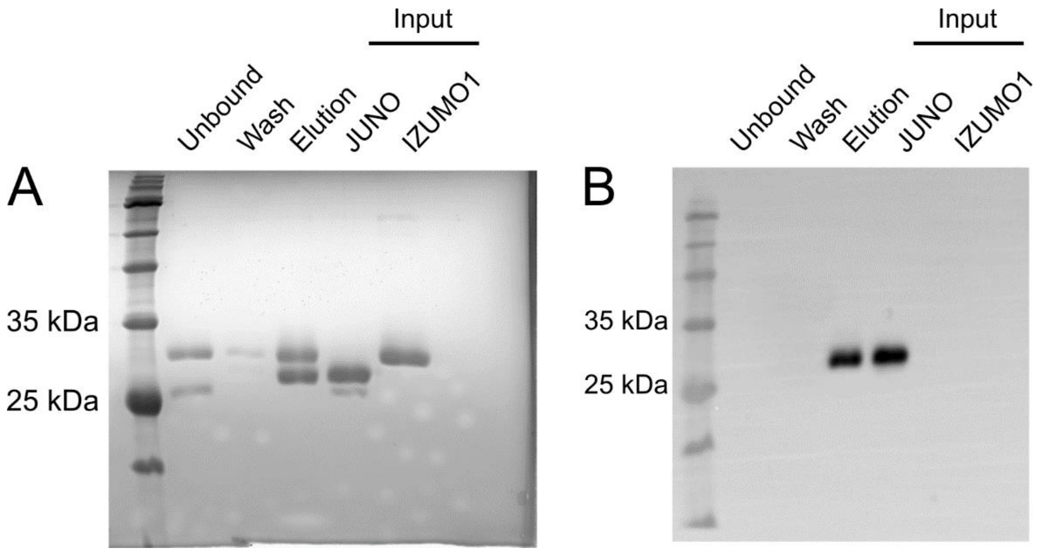

**Fig. S15. Positive control for pull-down assays. A)** Coomassie stained SDS-PAGE and **B)** Western blot of His-tagged JUNO (bait) incubated with untagged IZUMO1. Mouse anti-His primary antibody was used with an HRP-conjugated anti-mouse secondary for detection of proteins.

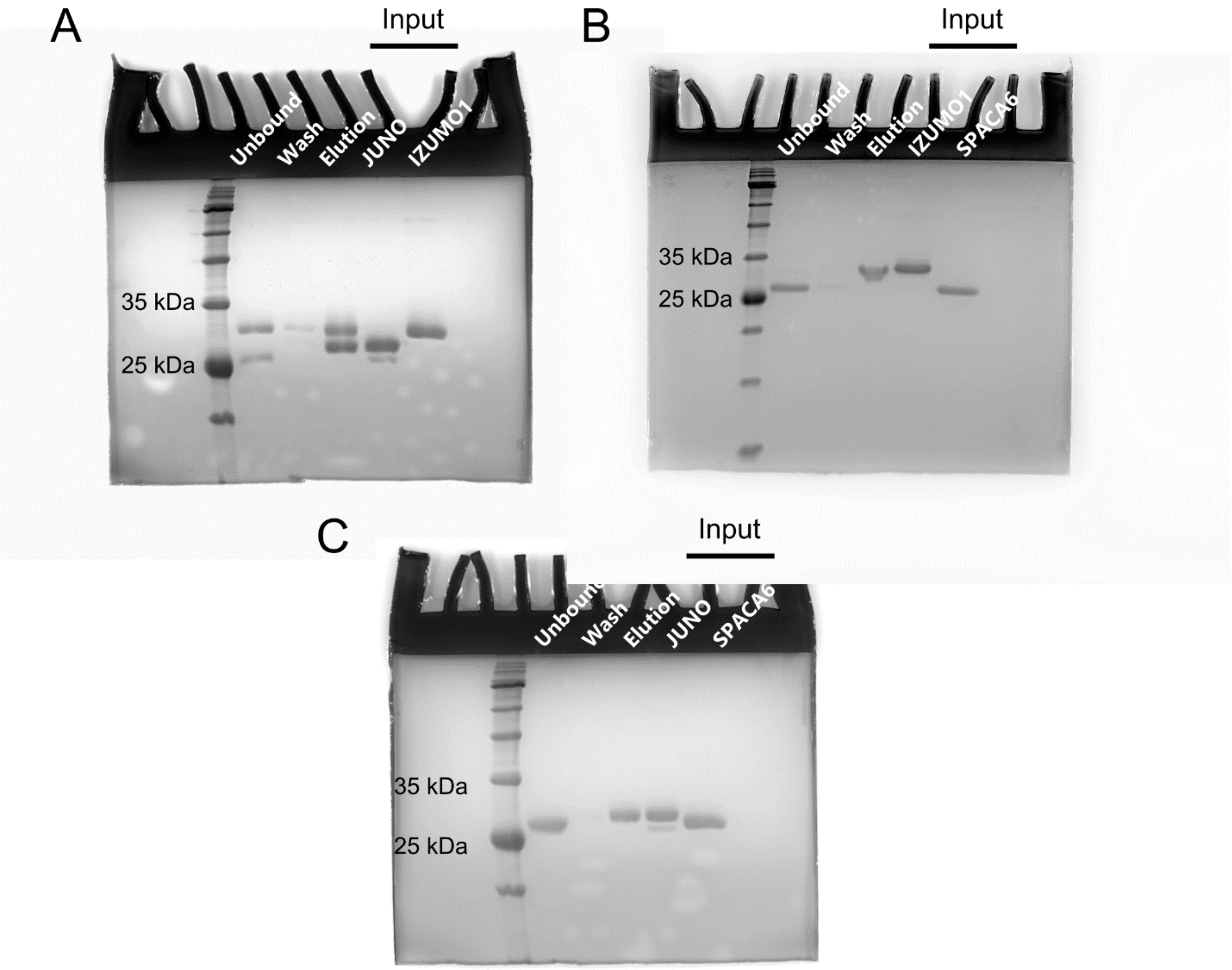

**Fig. S16. Source data for pull-down assays.** Uncropped Coomassie stained SDS-PAGE source data for: **A)** Supplementary Figure 15, **B)** Supplementary Figure 14C, and **C)** Supplementary Figure 14D.

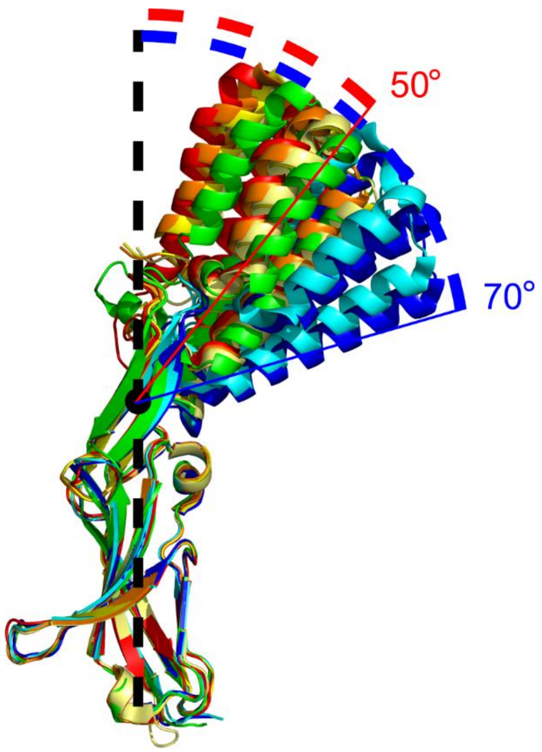

**Fig. S17. Crystallographic evidence of IZUMO1 flexibility.** Alignments of the multiple IZUMO1 solved structures by the Ig-like domain to compare orientation of the 4HB; the smallest and largest angles between the bottom of the Ig-like domain, the center of the hinge region, and the tip of helix 4 in the 4HB are displayed. IZUMO1 structures include PDB: 5F4E (red), 5JKC (orange), 5JKD (yellow), 5JK9 (light yellow), 5B5K (green), 5F4T (cyan), and 5F4V (blue).

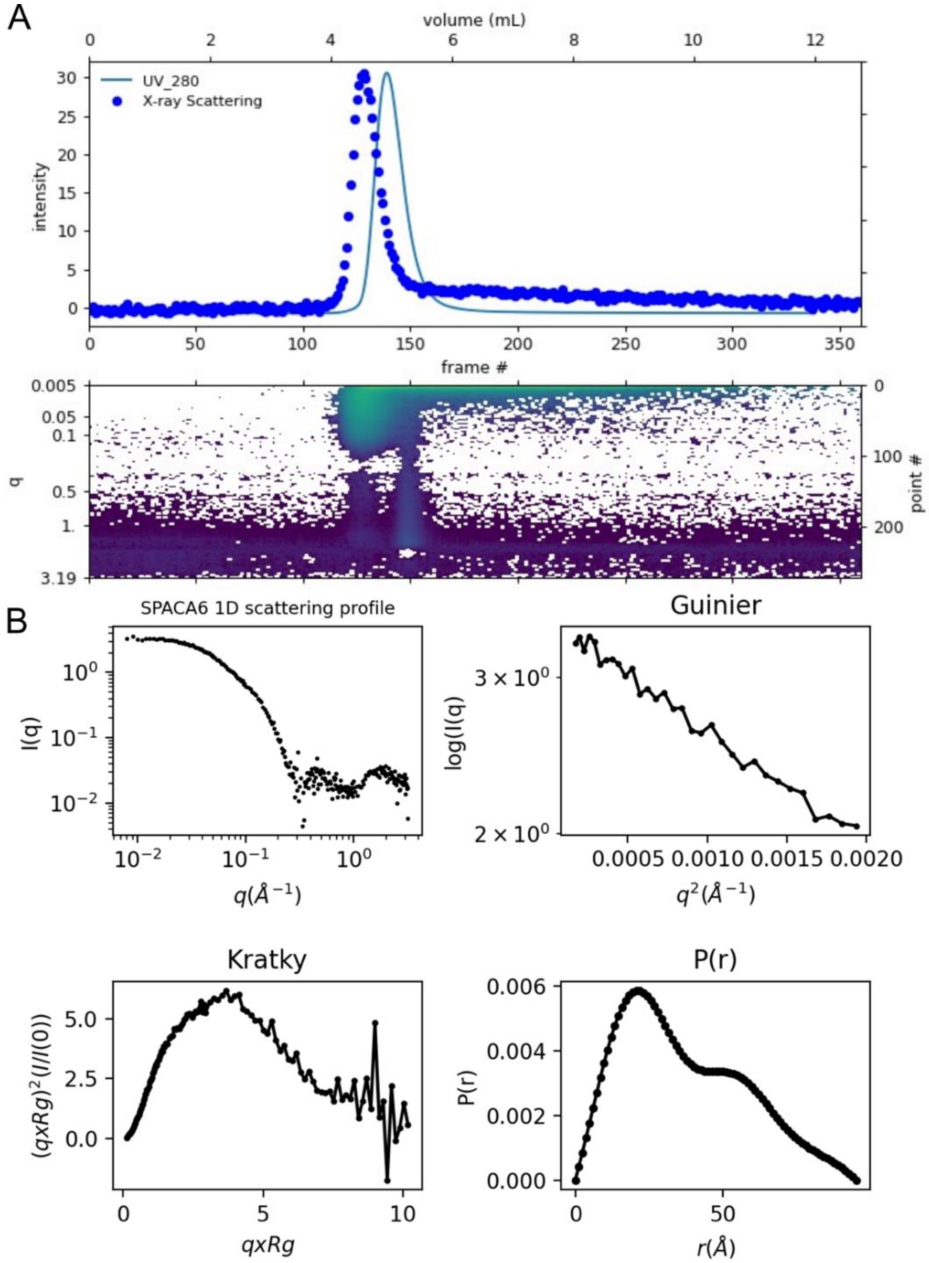

**Fig. S18. SEC-SAXS data of SPACA6. A)** Top panel shows X-ray scattering intensity (blue dots) and UV absorbance 280 nm trace (solid blue line) of SEC-SAXS elution profile (360 frames). Bottom panel represents a 2D heat map of the buffer subtracted X-ray intensity. Each vertical slice represents the X-ray intensity of the entire SAXS/WAXS q range from 0.005-3.1Å$^{-1}$ for each frame. **B)** Average intensity of frames 90-100 was used to perform buffer subtraction on frames 122-125 of the SPACA6 elution peak. A $R_g$ of 29.5 Å was calculated from the Guinier plot and $D_{max}$ of 95 Å was determined from the Distance Distribution function (P(r)). Dimensionless Kratky plot indicates the protein is well folded but displays flexibility.

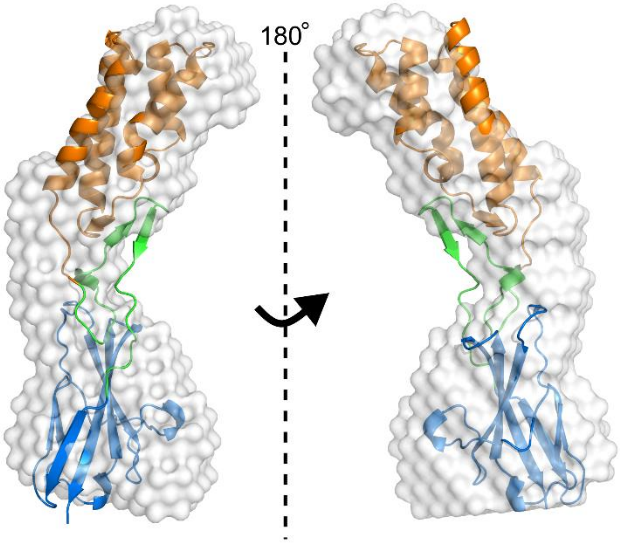

**Fig. S19. SAXS reconstruction of SPACA6.** *Ab initio* reconstruction of SPACA6 ectodomain using SAXS date (white envelope) overlaid with the SPACA6 crystal structure, coloured according to its three regions (4HB, orange; Ig-like domain, blue; hinge, green). DAMMIF was used to produce 20 envelope models, which were subsequently averaged with DAMAVER and filtered with DAMFILT.

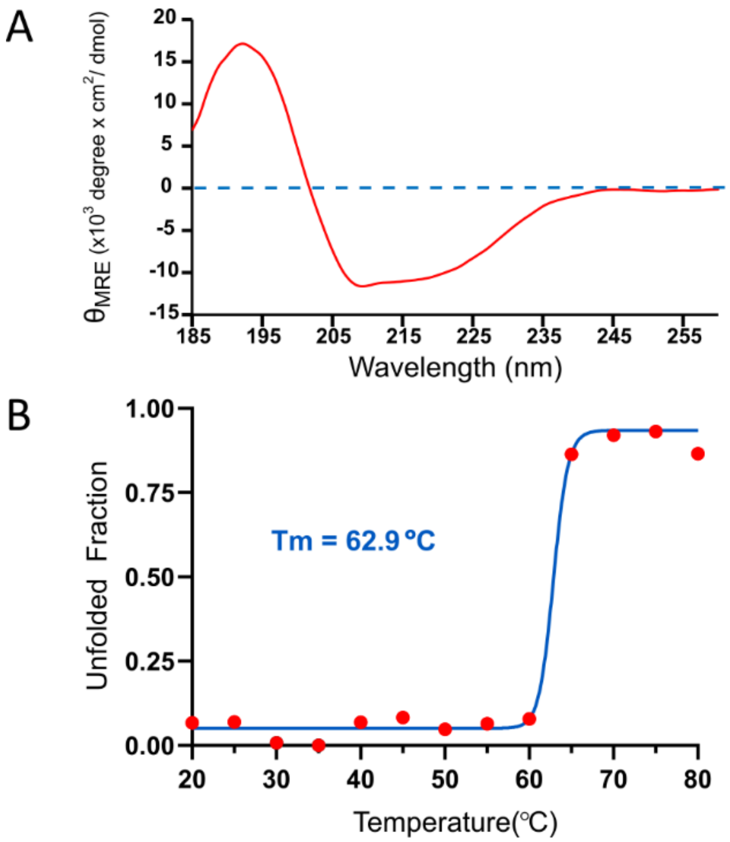

**Fig. S20. CD spectra and thermal melt of IZUMO1 ectodomain. A)** Far-UV CD spectra of the IZUMO1 ectodomain at 0.2 mg $mL^{-1}$. **B)** Thermal melt of IZUMO1 ectodomain in 10 mM sodium phosphate, pH 7.4, and 150 mM NaF.

**Table S1. Data collection and refinement statistics.**

| | ARCIMBOLDO SPACA6 | SPACA6 Br-SAD |
|---|---|---|
| Synchrotron | APS 24ID-C (NE-CAT) | APS 24ID-C (NE-CAT) |
| Wavelength (Å) | 1.6314 | 0.91165 |
| Resolution range (Å)[a] | 43.44 – 2.00 (2.072 – 2.000) | 44.83 – 2.25 (2.331– 2.25) |
| Space group | $P2_12_12_1$ | $P2_12_12_1$ |
| Unit cell<br>a, b, c (Å)<br>α, β, γ (°) | 27.6 44.5 195.9<br>90 90 90 | 29.0 46.5 167.7<br>90 90 90 |
| Total reflections[a] | 57321 (3492) | 82666 (6844) |
| Unique reflections[a] | 16683 (1503) | 11,435 (1107) |
| Multiplicity[a] | 3.4 (3.2) | 7.2 (6.7) |
| Completeness (%)[a] | 96.7 (90.1) | 99.8 (99.9) |
| Mean I/sigma(I)[a] | 8.6 (1.2) | 11.7 (2.8) |
| Wilson B-factor | 37.82 | 34.32 |
| Rmerge (%)[a,b] | 6.6 (65.4) | 10.8 (71.6) |
| Rmeas (%)[a,c] | 8.8 (91.8) | 11.6 (81.5) |
| Rpim (%)[a,d] | 4.6 (48.5) | 4.1 (30.3) |
| CC1/2[a] | 0.997 (0.675) | 0.998 (0.871) |
| R-work (%)[a,e] | 27.6 (29.3) | 20.1 (22.0) |
| R-free (%)[a] | 29.8 (30.5)[f] | 25.5 (30.1)[g] |
| Number of non-hydrogen atoms | 1592 | 1811 |
| macromolecules | 1592 | 1736 |
| bromide | N/A | 10 |
| solvent | 0 | 65 |
| RMSD (bonds; Å) | 0.01 | 0.005 |
| RMSD (angles; °) | 1.20 | 1.02 |
| Ramachandran favored (%) | 90.3 | 95.41 |
| Ramachandran allowed (%) | 6.6 | 4.59 |
| Ramachandran outliers (%) | 3.1 | 0.0 |
| Clashscore | 9.59 | 1.76 |
| Average B-factor ($Å^2$) | 44.5 | 44.0 |
| macromolecules | 44.5 | 44.31 |
| bromide | N/A | 50.86 |
| solvent | N/A | 34.41 |
| Number of TLS groups | N/A | 7 |
| PDB accession number | | 7TA2 |

[a]Statistics for the highest-resolution shell are shown in parentheses.
[b]$R_{merge} = \Sigma_{hkl}\Sigma_j|I_j - \langle I\rangle|/\Sigma_{hkl}\Sigma_j|\langle I\rangle|$, where $I_j$ and $\langle I\rangle$ represent the diffraction intensity values of the individual measurements and the corresponding mean values, respectively. The summation is over all unique measurements.
[c]$R_{meas} = \Sigma_{hkl}\sqrt{(n-(n-1))}\Sigma_j|I_j - \langle I\rangle|/\Sigma_{hkl}\Sigma_i|\langle I\rangle|$, where n is the number of diffraction intensities summated.
[d]$R_{pim} = \Sigma_{hkl}\sqrt{(1/(n-1))}\Sigma_j|I_j - \langle I\rangle|/\Sigma_{hkl}\Sigma_i|\langle I\rangle|$.
[e]$R_{work}$: = $\Sigma|\ |F_{obs}|-|F_{calc}|\ |\ /\ \Sigma|F_{obs}|$, where $F_{calc}$ and $F_{obs}$ are the calculated and observed structure factor amplitudes, respectively
$R_{free}$: statistic is the same as $R_{work}$ except calculated on [f]5.4% and [g]5.1% of the total unique reflections chosen randomly and omitted from the refinement

**Table S2. Dali server search of PDB.**

| Protein Name | PDB | Z-score | RMSD | LALI | %ID |
|---|---|---|---|---|---|
| **Full Ectodomain*** | | | | | |
| IZUMO SPERM-EGG FUSION PROTEIN 1 | 5jk9-A | 12.1 | 3.9 | 130 | 15 |
| DOWN SYNDROME CELL ADHESION MOLECULE | 3dmk-A | 9.2 | 6.7 | 92 | 13 |
| MUCOSA-ASSOCIATED LYMPHOID TISSUE TRANSLOCATION PROTEIN | 3bfo-A | 8.8 | 1.7 | 75 | 15 |
| COXSACKIE AND ADENOVIRUS RECEPTOR | 3j6n-K | 8.6 | 2.3 | 81 | 21 |
| CELL SURFACE GLYCOPROTEIN CD200 RECEPTOR 1 | 4bfi-A | 8.6 | 2.8 | 82 | 17 |
| LEUCINE-RICH REPEAT AND IMMUNOGLOBULIN-LIKE DOMAIN | 4oqt-A | 8.6 | 13.4 | 118 | 14 |
| INTERLEUKIN-33 | 5vi4-F | 8.5 | 3.7 | 97 | 9 |
| BASIC FIBROBLAST GROWTH FACTOR RECEPTOR 1 | 2cr3-A | 8.5 | 3 | 80 | 18 |
| CD2 | 1hng-A | 8.3 | 2.2 | 78 | 13 |
| ADVANCED GLYCOSYLATION END PRODUCT-SPECIFIC RECEPTOR | 4lp5-A | 8.2 | 2.7 | 80 | 15 |
| **Four-Helix Bundle†** | | | | | |
| IZUMO SPERM-EGG FUSION PROTEIN 1 | 5jk9-A | 7.2 | 3.5 | 116 | 8 |
| DESIGNED HELICAL REPEAT PROTEIN | 5cwc-A | 6.5 | 2.9 | 86 | 14 |
| PHOSPHOPROTEIN | 3l32-B | 6.2 | 1.2 | 43 | 5 |
| 55-KDA IMMEDIATE-EARLY PROTEIN 1 | 6tgz-F | 6 | 5 | 95 | 7 |
| DOLICHYL-DIPHOSPHOOLIGOSACCHARIDE PROTEIN | 6s7o-E | 5.7 | 3.7 | 66 | 9 |
| SOLUBLE CYTOCHROME B562 | 4or2-B | 5.5 | 4.5 | 68 | 7 |
| GLUTAMYL-TRNA REDUCTASE 1 | 5che-A | 5.5 | 6.7 | 62 | 3 |
| DNA-DEPENDENT RNA POLYMERASE SUBUNIT RPO132 | 6rfl-C | 5.5 | 7.5 | 56 | 4 |
| SPOROZOITE MICRONEME PROTEIN | 4u5a-B | 5.5 | 3.2 | 90 | 9 |
| RHUL123 | 4wid-A | 5.4 | 7 | 94 | 7 |
| **Ig-like domain‡** | | | | | |
| IZUMO SPERM-EGG FUSION PROTEIN 1 | 5jk9-A | 11.9 | 3.5 | 113 | 18 |
| DOWN SYNDROME CELL ADHESION MOLECULE | 3dmk-A | 9.2 | 4.4 | 89 | 12 |
| INTERFERON ALPHA-5 | 3oq3-B | 8.7 | 2.5 | 88 | 15 |
| COXSACKIE AND ADENOVIRUS RECEPTOR | 3j6n-K | 8.6 | 2.4 | 81 | 20 |
| MUCOSA-ASSOCIATED LYMPHOID TISSUE TRANSLOCATION PROTEIN | 3bfo-A | 8.6 | 1.7 | 74 | 15 |
| BASIC FIBROBLAST GROWTH FACTOR RECEPTOR 1 | 2cr3-A | 8.5 | 4.1 | 83 | 17 |
| LEUCINE-RICH REPEAT AND IMMUNOGLOBULIN-LIKE DOMAIN | 4oqt-A | 8.5 | 8.5 | 88 | 17 |
| CELL SURFACE GLYCOPROTEIN CD200 RECEPTOR 1 | 4bfi-A | 8.5 | 2.9 | 81 | 20 |
| TYROSINE-PROTEIN PHOSPHATASE | 4yh7-B | 8.5 | 2.5 | 89 | 18 |
| IP13724P | 6qp8-A | 8.3 | 2.7 | 101 | 11 |

* SPACA6 ectodomain structure, residues 27-246, was used to search the PDB25 subset of the Protein Data Bank.
† SPACA6 4HB + Hinge structure, residues 27-150, was used to search the PDB25 subset of the Protein Data Bank.
‡ SPACA6 Hinge + Ig-like domain structure, residues 121-246, was used to search the PDB25 subset of the Protein Data Bank.
